# Nodal Modulator (NOMO) is a force-bearing transmembrane protein required for muscle differentiation

**DOI:** 10.1101/2024.09.06.611727

**Authors:** Brigitte S Naughton, Swapnil C Devarkar, Sunanda Mallik, Stacey Oxendine, Sanjana Junnarkar, Yuan Ren, Vanessa Todorow, Julien Berro, Janine Kirstein, Yong Xiong, Christian Schlieker

## Abstract

The endoplasmic reticulum (ER) relies on specialized membrane-shaping proteins to maintain a continuous network of sheets and tubules that host distinct biological processes. How this intricate structure of the ER membrane system is maintained under conditions of mechanical strain is incompletely understood. NOMO is an ER-resident transmembrane protein that contributes to ER morphology and is highly expressed in striated muscle. In this study, we identify a critical interface between distal Ig domains that enables NOMO to maintain ER morphology and buffer mechanical forces. By incorporating two independent tension sensors in the luminal domain of NOMO, we demonstrate that NOMO assemblies experience forces in the single piconewton (pN) range, with a significant contribution from the identified interface. These newly defined features are important, if not indispensable, for myogenesis, as interface mutations affecting mechanosensing fail to restore the essential role of NOMO during myogenesis in a C2C12 differentiation model. Moreover, NOMO depletion impairs nematode motility, underscoring a broader functional importance in muscle physiology.

## Introduction

Nodal Modulator (NOMO) (also known as pM5) encodes a highly conserved type I transmembrane protein implicated in vertebrate development and endoplasmic reticulum (ER) structural integrity (Amaya et al., 2021; Haffner et al., 2004; Zhang et al., 2015; Zhao et al., 2017). While only one copy exists in other organisms, three nearly identical paralogs (NOMO1, NOMO2, NOMO3; hereafter referred to as NOMO) are encoded in the human genome, each featuring a large ER luminal domain, a single transmembrane domain, and a short cytosolic tail, with a total mass of ∼139 kDa. NOMO was first studied in zebrafish for its role in attenuating nodal signaling during embryonic cellular differentiation (Haffner et al., 2004; Shen, 2007). It associates with a variety of ER-resident proteins or protein complexes including Nicalin (Haffner et al., 2004), which interacts with TMEM147 (Dettmer et al., 2010), a resident of the back of Sec61 (BOS) complex implicated in multi-pass membrane protein biogenesis (Page et al., 2023; Sundaram et al., 2022). Ectopic expression of NOMO1 and Nicalin in zebrafish embryos induces cyclopia (Haffner et al., 2004) and NOMO is significantly downregulated in patients with facial asymmetry associated with skeletal malocclusion (Nicot et al., 2014). NOMO is highly expressed in heart and skeletal muscle (Uhlen et al., 2015) and reduced in patients with ventricular septal defect (VSD) (Zhang et al., 2006), a common congenital heart anomaly, signaling a contribution to cardiovascular health and disease.

These findings collectively suggest that NOMO plays a multifaceted role in both developmental and disease processes. Nevertheless, the cellular and molecular mechanisms underlying these observations are poorly understood. Further research into NOMO functional pathways is essential to elucidate disease etiology and potential therapeutic targets. In our previous work, we reported that NOMO depletion causes large fenestrations in the ER, whereas overexpression constricts ER sheets to an intermembrane spacing that scales with the molecular dimensions of NOMO1’s luminal domain (Amaya et al., 2021). However, the functional relevance in relation to molecular features of NOMO, particularly in muscle tissue and in an organismal context, remained unexplored.

In this study, we integrate Alphafold predictions and structural comparisons to identify an interface between distal NOMO Ig folds that we validate through *in vitro* reconstitution and determine to have modest affinity. These low-affinity interfaces are critical for the establishment of metastable NOMO assemblies that exhibit unusually slow diffusion dynamics within the ER. By applying two independent tension sensors, we demonstrate that NOMO assemblies experience intraluminal forces in the pN range. The force-bearing features identified in this study are crucial for NOMO’s mechanosensitive properties during myogenesis, as shown by the necessity of interface-proficient NOMO1 for myocyte differentiation in a C2C12 model. Lastly, silencing of the NOMO homolog *nra-4* in *Caenorhabditis elegans* significantly reduces the motility of the nematode. Collectively, our observations suggest an unexpected role for NOMO as an ER-resident mechanosensor, with implications for understanding NOMO-linked developmental disorders affecting skeletal muscle and heart function.

## Results

### NOMO comprises conserved Ig-like domains similar to those of force-bearing proteins

To gain insights into NOMO’s architecture, we previously purified and subjected full-length (FL) NOMO1-FLAG to negative-stain EM, which revealed a flexible, extended rod-like structure approximately 27 nm in length (Amaya et al., 2021). Two-dimensional classifications resolved a “beads on a string” morphology with eight discernible globular segments, possibly representing immunoglobulin (Ig)-like domains. However, the flexibility of NOMO posed experimental obstacles to resolving the full-length structure. High-confidence Alphafold3 predictions (pTM = 0.54) (Abramson et al., 2024) of NOMO (Fig. 1a-d and Extended Data Fig. 1a-d) have since corroborated the presence of twelve Ig-like domains that reside in the ER lumen, followed by a single transmembrane domain and a short cytosolic tail (Fig. 1a). The model posits NOMO as adopting a looping topology facilitated by an interface formed by distal Ig domains. Within this interface, we identified several salt bridge interactions involving Ig 1 D70 - Ig 10 K927, Ig 1 D121 – Ig 10 R903, and Ig 11 D994,E993 – Ig 1 K66, Ig 10 K931 (Fig. 1b,c). Consistent with an important role for this interface, residues contributing to this interface are evolutionarily conserved (Fig. 1d,e and Extended Data Fig. 1c) (Ashkenazy et al., 2016) (Waterhouse et al., 2009).

**Fig. 1.**
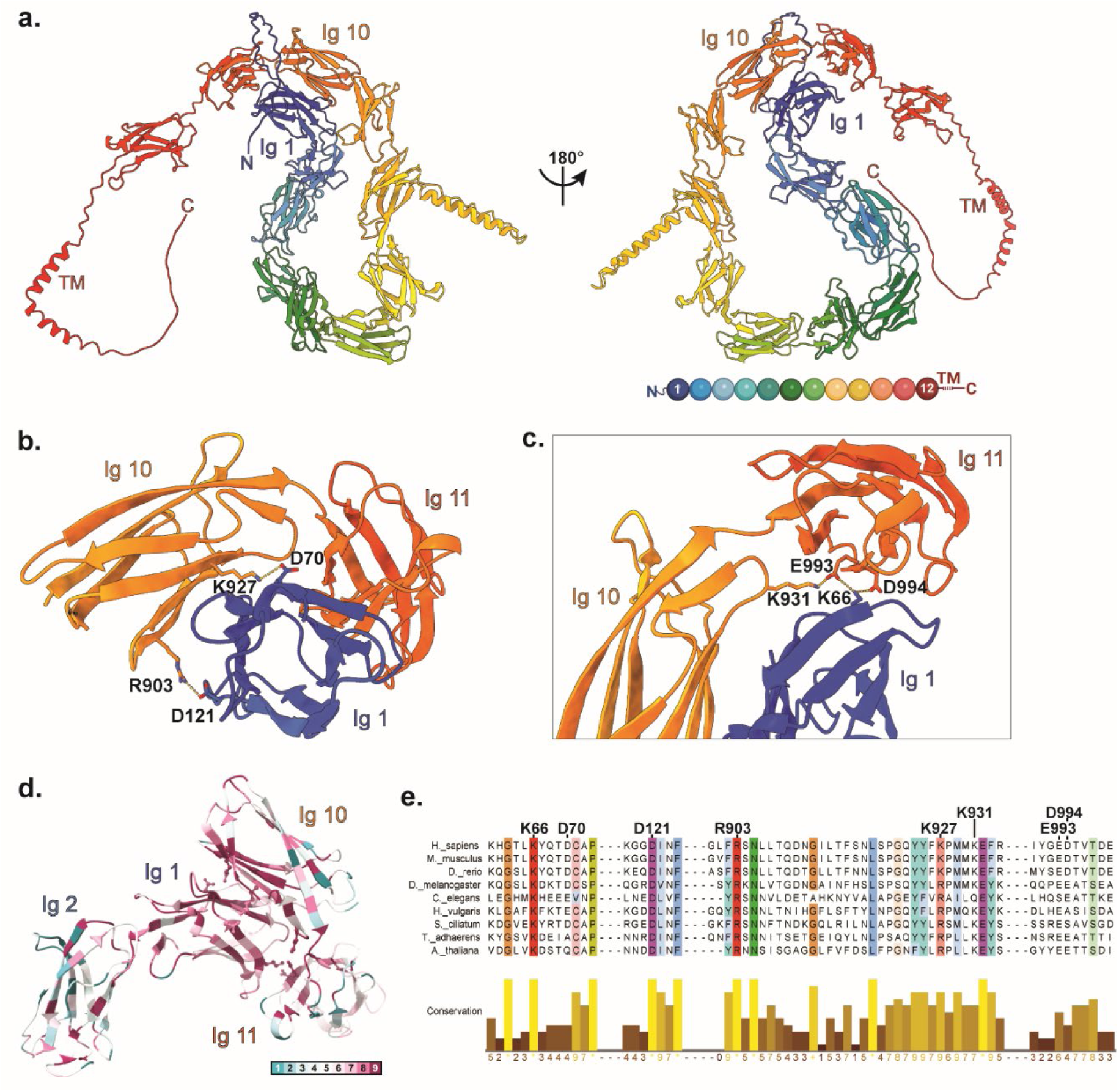
Predicted structural features of NOMO. a, An Alphafold3 model prediction of full-length human NOMO1 resolves twelve immunoglobulin-like (Ig-like) folds, colored by Ig domains depicted as balls in the bottom right cartoon. NOMO1 is represented in two views related by a 180° rotation, as indicated. b, Magnification of the interface formed by Ig 1/10/11 with D70-K927 and D121-R903 salt bridges labeled. c, An alternative view of the Ig 1/10/11 interface with K66-D994-K931-E993 salt bridges labeled. d, Ig 1-2 and Ig 10-11 interface conservation. Inset, ConSurf color-coding scheme from (1) variable to (9) conserved. e, Sequence conservation of regions around and including the salt bridges shown in b-c, colored by >80% conservation.

### Salt-bridge interactions are critical for NOMO function and bestow slow diffusional mobility on the NOMO assembly

The ER of wild-type U2OS cells forms an extensive and intricate network of tubules and sheets that spread throughout the cell, with reticular tubules predominantly in the peripheral regions and dense sheets predominantly near the nucleus (Fig. 2a, left). Upon NOMO depletion, membrane-delineated voids appear in the ER network, ranging from < 0.5 µm to 5 µm (Fig. 2a, right). Overexpression of an siRNA-resistant wild-type (WT) NOMO1 construct with an N-terminal Flag tag (F-NOMOr) rescues this phenotype (Fig. 2b, lower left, and Fig. 2d, green bar).

**Fig. 2.**
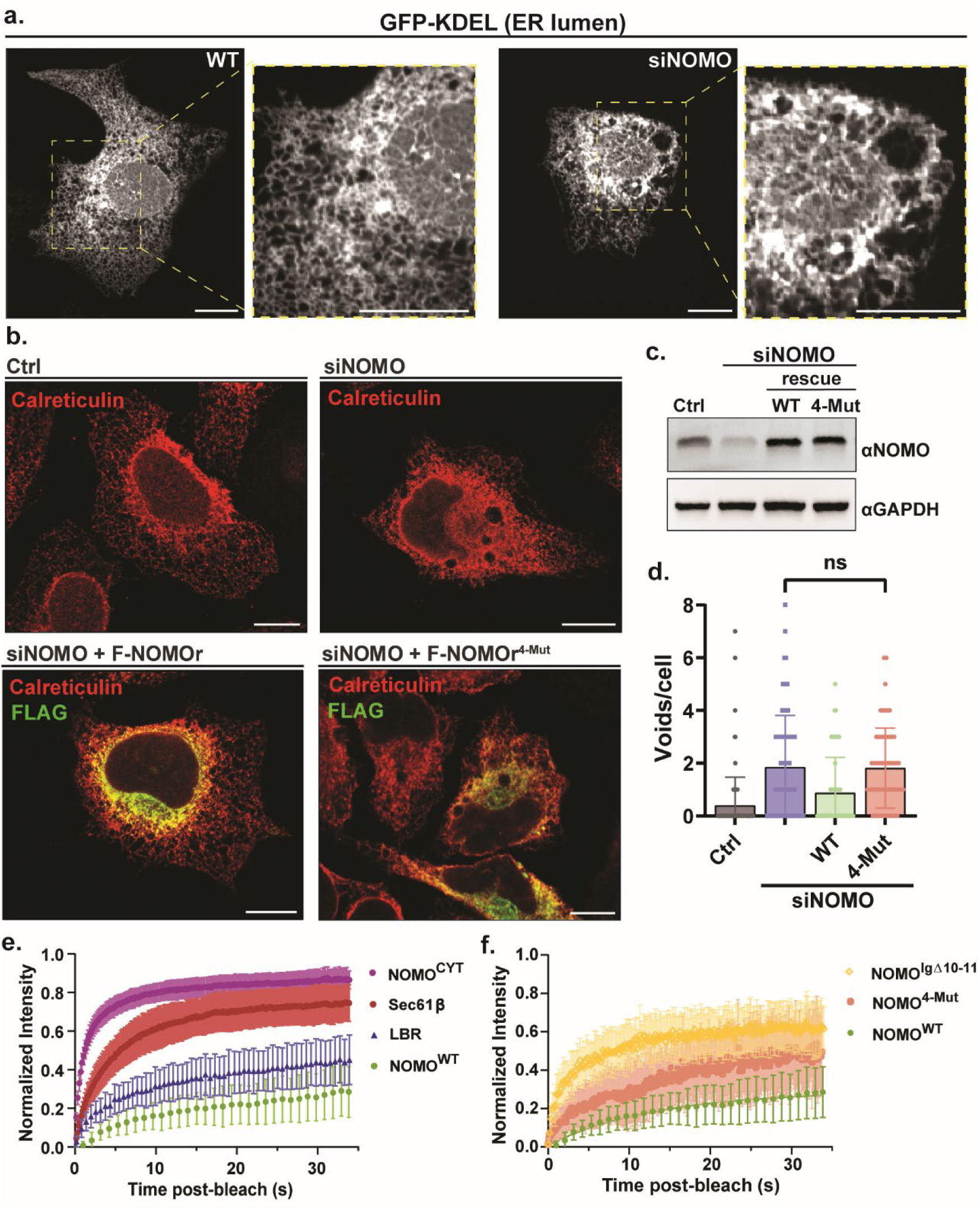
NOMO relies on the predicted Ig 1/10/11 interface for functionality. a, Spinning disc confocal images of U2OS cells under either untreated or NOMO siRNA-treated samples. b, Immunofluorescence images of U2OS cells treated with either non-targeting siRNA (Ctrl) or siRNA against NOMO (siNOMO) and transfected with constructs indicated in bottom two panels. c, Immunoblot of cell lysates from the transfection in b. d, Plot of ER voids per cell under indicated conditions. Statistical analyses were performed using a two-tailed unpaired Mann– Whitney test; \*\*\**P* < 0.001 and ns, not significant. e,f, FRAP data of means ± s.e.m. of fluorescence intensities of bleached regions normalized to the prebleach intensity for each condition. NOMO^WT^ data is the same in e,f. Scale bars, 10 μm. FRAP, fluorescence recovery after photobleaching.

To assess whether the Ig 1 and Ig 10-11 interface (Ig 1/10/11) is important for NOMO function, we engineered an siRNA-resistant NOMO1 construct with four charge-reversal mutations predicted to disrupt the interface (K66E, D70K, D121K, K931E, designated F-NOMOr^4-Mut^). We found that expression of F-NOMOr^4-Mut^ in a NOMO knockdown background did not rescue the void phenotype (Fig. 2b, lower right, and Fig. 2d, peach bar). Immunoblotting of cell lysates from each condition confirmed efficient depletion of endogenous NOMO and similar expression of both F-NOMOr and F-NOMOr^4-Mut^ (Fig. 2c). We conclude that the Ig 1/10/11 interface is important for sustaining ER morphology.

We next examined NOMO dynamics in the ER by employing fluorescence recovery after photobleaching (FRAP) in HeLa cells. ER-resident Sec61β with a GFP tag (Sec61β -GFP) served as a highly diffusive control (Shibata et al., 2008) and nuclear-resident GFP-tagged lamin B receptor (GFP-LBR) as a slowly diffusive control (Fig. 2e). F-NOMO1r was tagged with an internal eGFP-tagRFP moiety (TS) between Ig 12 and the TM domain (NOMO-TS_in_), explained in detail below. We additionally included a TS-tagged soluble C-terminal tail of NOMO (CYT) with rapid recovery kinetics to ensure the TS moiety did not interfere with mobility. We photobleached GFP with a 1 µm diameter ROI in either the ER (for Sec61β and NOMO) or nucleus (for LBR) and monitored recovery over time. In contrast to the rapid diffusion of Sec61β, NOMO exhibited markedly slow and restricted lateral mobility (Fig. 2e,g). Notably, NOMO’s recovery profile was even slower than that of LBR, which displays limited mobility due to its attachment to the nuclear lamina (Ellenberg et al., 1997) (Ungricht et al., 2015).

To test whether the Ig 1/10/11 influences NOMO’s diffusional dynamics, we transfected F-NOMO^4-Mut^ and a construct with Ig 10-11 removed via an in-frame deletion (F-NOMO^Δ10-11^). Each NOMO variant harbored an internal TS in the same position as the WT NOMO construct used in FRAP. Removing either the entire ∼20 kD region corresponding to Ig 10-11 or introducing the four point mutations at the Ig 1 and Ig 10-11 interface, increased NOMO mobility (Fig. 2f). Taken together, these FRAP data indicate that NOMO diffuses slowly within the ER membrane, relying in part on the presence of the Ig 1/10/11 interface.

### Ig 1-2 and Ig 10-11 dimerize *in vitro* and their expression induces a dominant-negative void phenotype

The inability of NOMO^4-Mut^ to rescue NOMO-depleted ER morphology prompted us to validate the Ig 1 and Ig 10-11 interaction with purified components. To this end, we individually expressed and purified recombinant His_6_-Ig 1-2 and His_6_-Ig 10-11 for *in vitro* binding assays (Fig. 3a). Due to the presence of a central beta sheet spanning Ig 1 and Ig 2, we purified a tandem construct to avoid structural perturbations. AlphaFold-Multimer generates a high-confidence model and interface between Ig 1-2 and Ig 10-11 (pTM = 0.85 and ipTM = 0.81, respectively) (Extended Data Fig. 1b). We turned to size-exclusion chromatography (SEC), incubating His_6_-Ig 1-2 with His_6_-Ig 10-11 for one hour before loading the mixture onto a Superdex 75 pg column. Compared to the individual SEC elution profiles of His_6_-Ig 1-2 and His_6_-Ig 10-11, the mixture exhibited a shift to a higher molecular weight, consistent with formation of a dimer (Fig. 3b, n=3). Introducing three point mutations at the salt bridge interface in Ig 1 (K66E, D70K, D121K) (His_6_-Ig 1-2^3-Mut^) eliminated the observed shift (Fig. 3c). To validate these results and quantify the interaction strength, we next performed isothermal calorimetry (ITC). Employing the same His_6_-Ig 1-2 and His_6_-Ig 10-11 fragments, we observed dimerization with a moderate affinity of 1.98 µM +/- 0.81 (Fig. 3d, n=3). In agreement with SEC results, we found no interaction between His_6_-Ig 1-2^3-Mut^ and His_6_-Ig 10-11 in ITC at the tested concentrations (Fig. 3e, n=3).

**Fig. 3.**
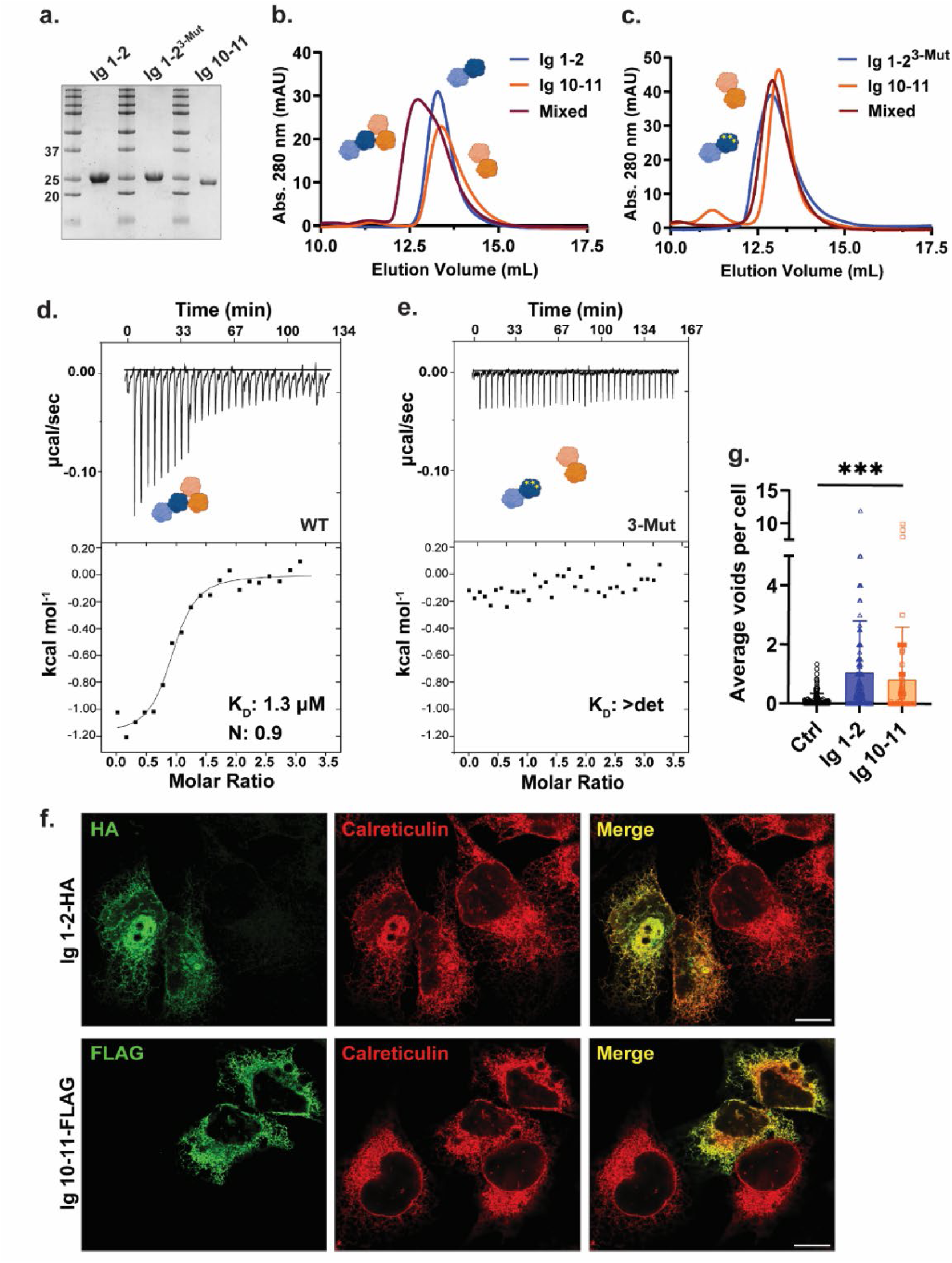
Ig 1-2 and Ig 10-11 dimerize and induce ER voids upon overexpression. a, SDS-PAGE/Coomassie staining of constructs obtained from preparative SEC. b,c, SEC profiles of recombinantly purified Ig 1-2 and Ig 10-11 on a HiLoad Superdex 75 pg. d,e, ITC binding studies of constructs used in b,c with dissociation constants (K_D_) and binding stoichiometry (n). f, U2OS cells immunostained for the ER marker calreticulin and either Ig 1-2-HA or Ig 10-11-FLAG. g, Plot of average voids per cell per image counted under indicated conditions. Error bars represent ± s.e.m. Statistical analyses were performed using a two-tailed unpaired Mann– Whitney test; ***P < 0.001. Scale bars, 10 μm. SEC, size-exclusion chromatography; ITC, isothermal calorimetry.

Considering that salt bridge interactions between Ig 1-2 and Ig 10-11 are integral to NOMO function (Fig. 2b), we next investigated whether these fragments could interfere with endogenous NOMO activity. To explore this, we individually expressed either Ig 1-2-HA or Ig 10-11-FLAG in WT U2OS cells and assessed ER morphology. Interestingly, expressing either construct induced a dominant-negative effect (Fig. 3f,g) reminiscent of NOMO depletion (Fig. 2a,b), though to a lesser extent. This suggests that the NOMO assembly is dynamic, as ectopically expressed Ig modules can compete for the same interface to perturb NOMO function. Together, these results demonstrate that a moderately strong interaction between Ig 1 and Ig 10-11 is critical for NOMO function.

### The Ig 1/10/11 interface maintains a compact NOMO topology

To evaluate how Ig 1-2 and Ig 10-11 interactions and ER disruption relate to the failure of NOMOr^4-Mut^ to rescue NOMO depletion, we employed SEC coupled with multiangle light scattering (SEC-MALS). In our previous analysis, we found that WT NOMO1-FLAG has a radius of gyration (Rg) of approximately 15 nm and a protein detergent complex (PDC)-corrected molar mass ranging from 230-270 kDa, indicative of dimer formation (Amaya et al., 2021). Consistent with this, we observed a molar mass ranging from 220-270 kDa and a Rg of 15 nm for WT NOMO1-FLAG (Fig. 4a). The experimental radius is similar to that predicted by Alphafold modeling of WT NOMO, which projects distances of 12-18 nm between distal points of the ER luminal domain in three-dimensional space (Extended Data Fig. 1d). NOMOr^4-Mut^-FLAG exhibited a comparable molar mass range to WT NOMO, suggesting it retains dimerization capability. However, it had a significantly greater Rg of approximately 40 nm (Fig. 4b). The twelve Ig domains modeled by Alphafold average approximately 3.5 nm per fold, or 42 nm cumulatively, which is compatible with an elongated conformation for the interface-deficient NOMO or a looping topology adopted by WT NOMO (Fig. 4c,d).

**Fig. 4.**
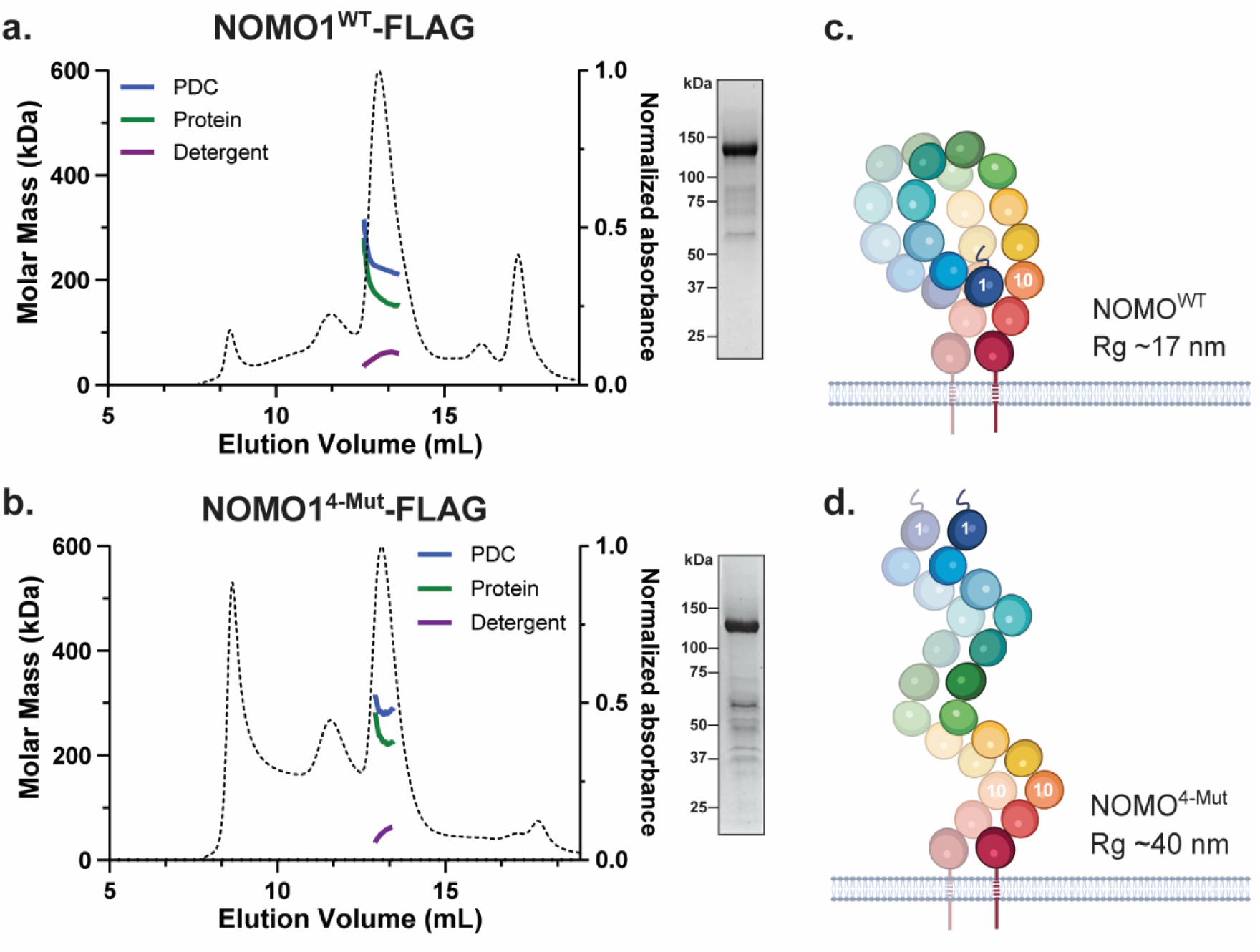
NOMO architecture is shaped by the Ig 1/10/11 interface. a, SEC-MALS profile of FL WT NOMO (NOMO1^WT^-FLAG) on a Superose 6 column. The dashed line is the UV trace, molar mass represents the total molar mass of the PDC, and protein molar mass is the corrected molar mass to remove contribution from detergent. Inset, SDS-PAGE/Coomassie staining of the NOMO1^WT^-FLAG fraction obtained from preparative SEC. b, Model representation based on the results in panel a. c, SEC-MALS profile of the 4-mutant interface NOMO1 mutant, NOMO^4Mut^-FLAG, insert as in a. d, Cartoon model representation based on the results in panel c. SEC-MALS, size-exclusion chromatography linked to multiangle light scattering.

### NOMO experiences interface-dependent force *in situ*

NOMO bears structural similarity to titin and several bacterial pili proteins with mechanosensitive Ig folds, which are known to unfold and refold in response to stress (Supplemental Table 1). These include BcpA, the major pilin subunit of *Bacillus cereus* (PDB ID 3KPT) (Budzik et al., 2009), RrgA and RrgC, pilus subunits of *Streptococcus pneumoniae* (PDB ID 2WW8 and 4OQ1) (Echelman et al., 2016; Shaik et al., 2014), and the minor pilin of *Streptococcus agalactiae* (PDB ID 3PHS) (Krishnan et al., 2007). In light of this, we sought to determine whether NOMO experiences force in the natural context of the ER. To do so, we implemented a cleavage-based readout that relies on TEV protease (TEVp) activity. This approach relies on insertion of a TEV recognition sequence into a linker region between a coiled-coil interface of known mechanical strength (Ren et al., 2023). In this constrained conformation, the TEV recognition sequence is too distorted to be cleavable by the protease unless the coiled-coil interface is disrupted by a defined force threshold, which exposes the TEV recognition sequence for proteolysis (Fig. 5a). The sensors were calibrated by optical tweezers that determined either 5 pN or 10 pN of force is required to unfold the coiled-coil and expose the cleavage site acted upon by co-transfected TEVp in cells. We placed either sensor between Ig 11 and Ig 12 (NOMO^WT^_11-cc-12_) and appended an N-terminal signal sequence (SS) and C-terminal KDEL sequence to TEVp (TEVp-HA) to direct it to the ER lumen of U2OS cells (Fig. 5b,c). Strikingly, the coiled-coil 5 pN sensor (cc-5 pN) was cleaved completely, while the coiled-coil 10 pN sensor (cc-10 pN) remained uncut (Fig. 5d), indicating that NOMO is under force in the ER in the single pN range. We next asked if the previously defined Ig 1/10/11 interface is involved in force transduction across the ER lumen. Indeed, installation of the 5 pN sensor into the interface mutant derivative (NOMO^4-Mut^_11-5pN-12_) resulted in approximately 50% reduction in cleavage, indicating that mechanical force experienced by NOMO depends partially on Ig 1/10/11.

**Fig. 5.**
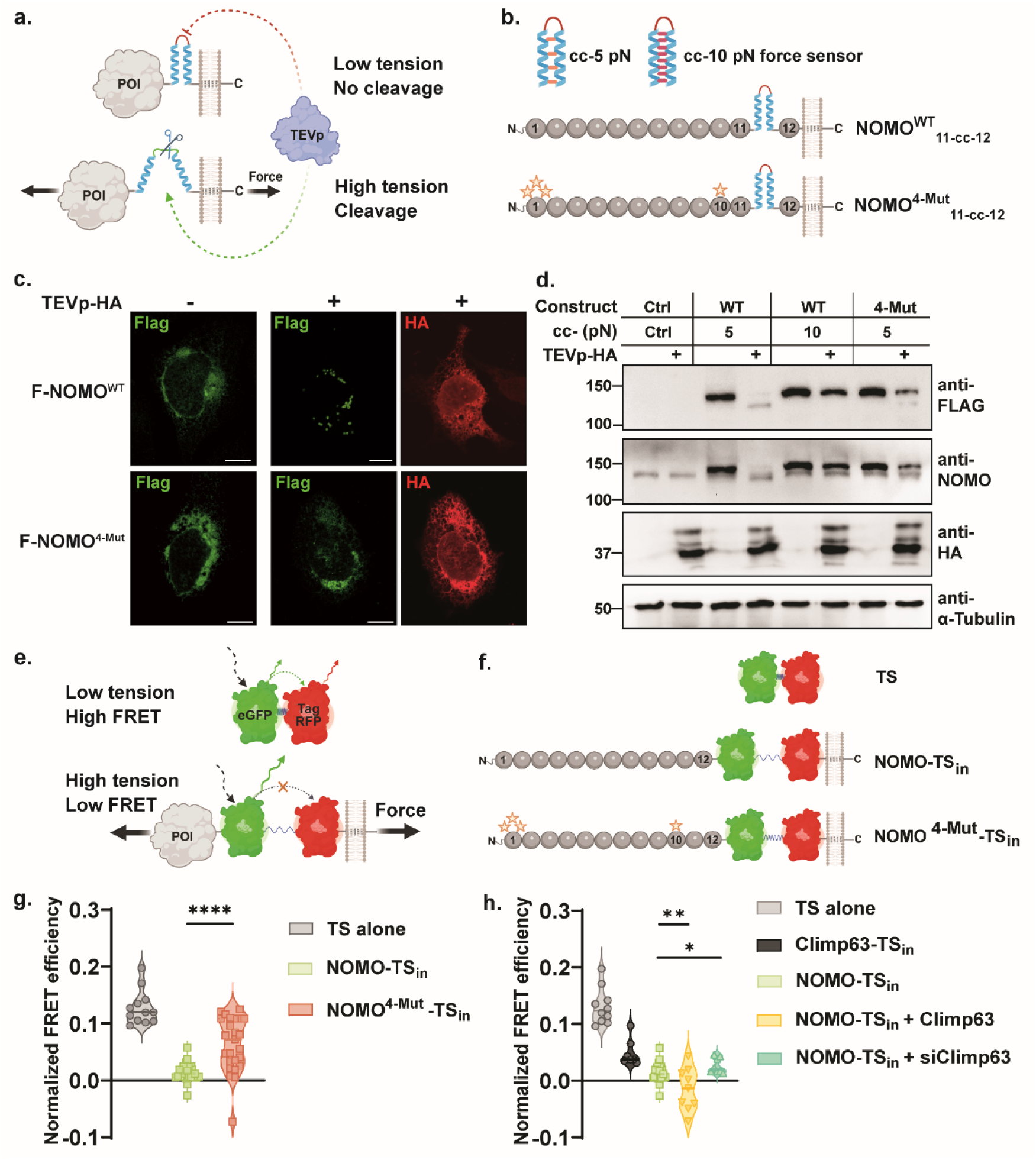
NOMO is under force dependent on the critical Ig 1/10/11 interface. a,b, Illustrations of the TEV cleavage-based readout and constructs used in c,d. c, Immunoblot of the cc-5 pN and cc-10 pN sensor in WT NOMO and the cc-5 pN sensor in NOMO^4-Mut^ when TEV protease (TEVp) is absent or present (+). d, Immunofluorescence images of conditions shown in c. of WT or 4-mutant NOMO in either the absence (-) or presence (+) of TEVp. e,f, Depictions of the FRET-based force sensor readout inserted between Ig 12 and the TM domain (TS_in_) of constructs used in g,h. g, Plots of normalized average FRET efficiency for WT (NOMO-TS_in_) and 4-mutant interface (NOMO^4-Mut^-TSin) constructs and a tension-sensor only control (TS). h, Normalized average FRET efficiency for TS, Climp-63 with TS inserted in the luminal domain prior to the TM domain (Climp-63-TS_in_), and NOMO-TS_in_ under control conditions (light green, as in g), with Climp-63 overexpression (yellow), or Climp-63 knockdown (aqua). Statistical analyses were performed using a two-tailed unpaired Mann–Whitney test; \*\*\*\**P* < 0.0001, \*\**P* < 0.01, \**P* < 0.05.

As an orthogonal and complementary approach, we employed a well-established FRET-based tension sensor (TS) (Kumar et al., 2016) that relies on an elastic flagelliform sensitive in the range of 1-6 pN (Grashoff et al., 2010). In its relaxed state, the nanospring is compact, resulting in high FRET; applied tension stretches the spring, reducing FRET intensity (Fig. 5e). The module was inserted after Ig 12 and before the TM domain in either WT Flag-NOMO1 (NOMO-TS_in_) or 4-mutant Flag-NOMO1 (NOMO^4-Mut^-TS_in_) (Fig. 5f). Consistent with the cleavage-based approach, WT NOMO was under high force (low FRET) while NOMO^4-Mut^ was under reduced force (moderate FRET) (Fig. 5g). We additionally installed the tension sensor in the ER spacer Climp-63, which experiences less force than NOMO (Fig. 5h). In order to test whether mechanical strain through NOMO could be modulated, we used Climp-63 to vary ER luminal width, as we previously observed genetic interactions between Climp-63 and NOMO in modulating ER morphology (Amaya et al., 2021). Overexpression of Climp-63 leads to an increase in both the abundance and luminal width of sheets (Shibata et al., 2010), which moderately elevated tension through NOMO (Fig. 5h). Conversely, depletion of Climp-63 narrows the luminal width of the ER (Shen et al., 2019), which resulted in reduced force on NOMO. Thus, we employed two independent readouts to arrive at the same conclusion: NOMO experiences force in the single pN range, with a critical contribution from the Ig 1/10/11 interface.

### NOMO is required and relies on the Ig 1/10/11 interface for myogenesis in a C2C12 model and depletion reduces nematode motility

Considering connections between NOMO and cardiogenesis (Xing et al., 2021; Zhang et al., 2015; Zhang et al., 2006), high expression in heart and muscle tissue (Uhlen et al., 2015), as well as load-bearing capabilities described above, we examined available transcriptomic data on muscular dystrophies. Interestingly, we identified differential NOMO1 expression in myotonic dystrophy type 1 (DM1) patients, with a positive correlation to dorsiflexion strength in the tibialis anterior muscle (R = −0.59) (Extended Data Fig. 4) (Wang et al., 2019). These findings prompted us to investigate whether NOMO and its mechanosensing properties play a role in myogenesis. Using the well-established mouse C2C12 *in vitro* differentiation model (Extended Data Fig. 2a), we performed an acute murine NOMO (_m_NOMO) KD (Fig. 6a,b), in addition to generating _m_NOMO KO myoblast lines (Extended Data Fig. 2b). _m_NOMO depletion minimally impacted myoblast survival and moderately delayed proliferation, arguing against nonspecific defects affecting viability. In differentiating myoblasts, cells fuse into multinucleated myotubes and express the differentiation marker Type I myosin heavy chain (Myhc) (Fig. 6a, and Extended Data Fig. 2a-c), among other protein indicators (Chal & Pourquie, 2017). Upon inducing myogenesis, however, _m_NOMO-depleted cells exhibited significantly delayed and incomplete differentiation (Fig. 6a,b, lower images). Despite expressing comparable levels of Myhc to WT cells (Extended Data Fig. 2c), _m_NOMO knockdown resulted in irregular clusters of Myhc rather than a uniform distribution along the myotubes, similar to the phenotype reported upon depletion of the fusion-promoting protein myomaker (Millay et al., 2013). Moreover, these cells formed thinner tubules containing fewer nuclei (Fig. 6a,d). In _m_NOMO KO lines, failure to express Myhc may be attributable in part to repeated passaging to isolate clones, likely exacerbating myogenesis impairment by NOMO depletion (Extended Data Fig. 2b,e,f) (VanGenderen et al., 2022). However, _m_NOMO depletion via siRNA knockdown is highly efficient and can be used as an alternative to avoid prolonged passaging (Extended Data Fig. 2c). We additionally probed for effects to ER morphology and actin organization in myoblast lines lacking _m_NOMO (Extended Data Fig. 3 a,b). Upon NOMO silencing, ER morphology was moderately perturbed in undifferentiated myoblasts, while actin organization was irregular in both undifferentiated (Day 0) and differentiating myotubes (Day 3). Overall, _m_NOMO depletion led to disrupted Myhc organization and ectopic clustering upon RNAi (see arrowheads, Fig. 6a) and an apparent lack of Myhc expression in KO cells, ultimately resulting in failure to establish myotubes in both conditions (Fig. 6a,d and Extended Data Fig. 2b,f). This was also evident in EM micrographs, which additionally revealed a nuclear envelope (NE) deformation phenotype upon NOMO depletion (Fig. 6b, lower).

**Fig. 6.**
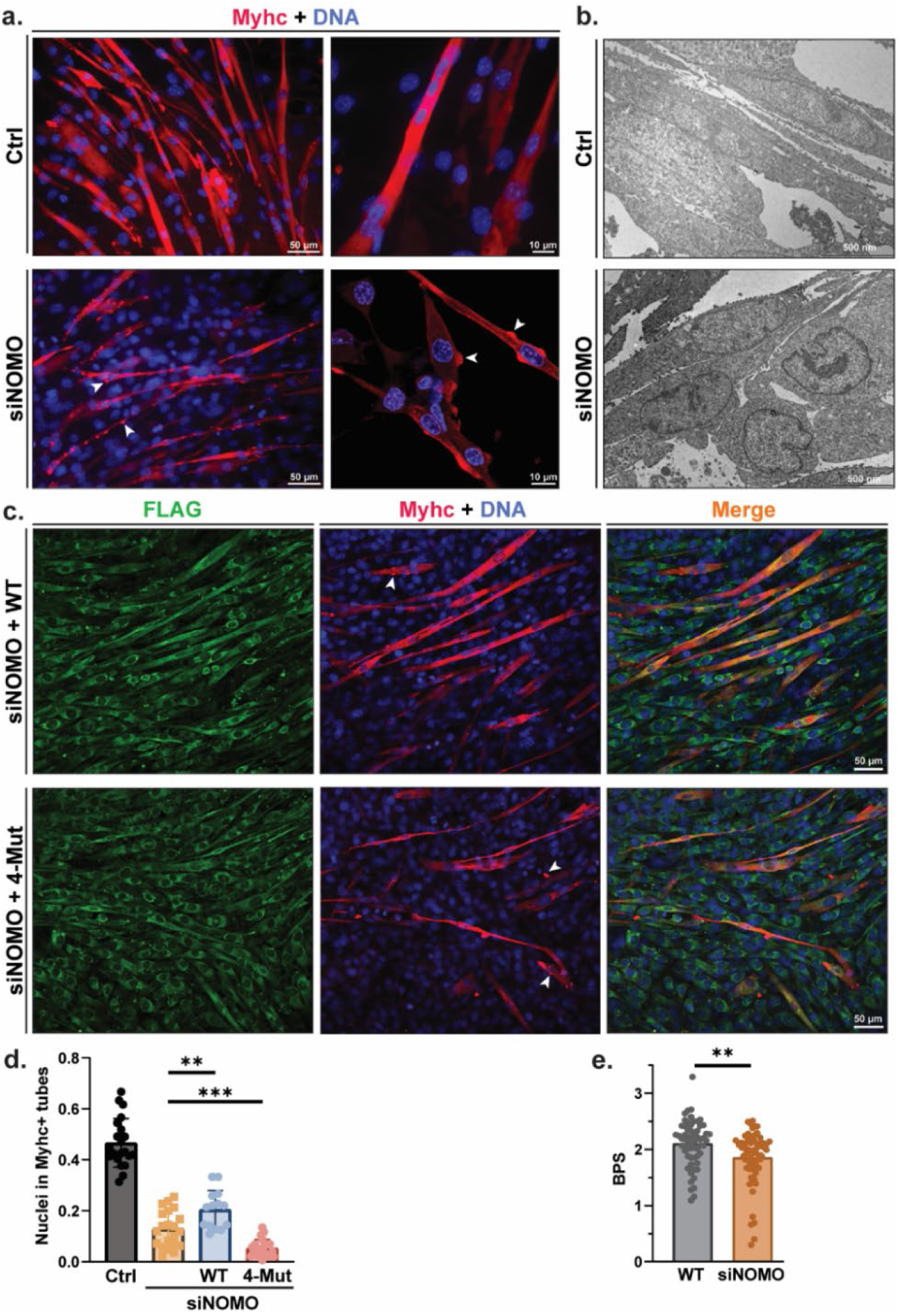
NOMO is required for myogenesis and depends on the Ig 1/10/11 interface. a, Cells fixed and immunostained for Myhc and stained for DAPI (DNA) three days post differentiation induction of myoblasts; left scale bars, 50 μm; right scale bars, 10 μm. b, Electron micrograph of either non-targeting (Ctrl) or NOMO knockdown (siNOMO) myoblasts on day three of differentiation. c, Assay performed as in a, with myoblasts stably expressing siRNA-resistant FLAG-NOMO1r (WT) or FLAG-NOMO1r^4-Mut^ (4-Mut) constructs under NOMO knockdown. d, Myogenesis quantification, scored by the fraction of nuclei in myosin heavy chain positive (Myhc+) myotubes. e, Motility analysis measured by bends per second (BPS) of *C. elegans* challenged with either non-targeting (n = 58) or *nra-4*-directed RNAi (n = 64). Data shows ± s.e.m. Statistical analyses were performed using a two-tailed unpaired Mann–Whitney test; ***P < 0.001, **P<0.01.

To verify the specificity of _m_NOMO depletion, we conducted NOMO rescue experiments in _m_NOMO KD and KO backgrounds. C2C12 lines were engineered to stably express WT human F-NOMO1r at moderate levels, as confirmed by IF and immunoblot (Fig. 6c and Extended Data Fig. 2d,e). In _m_NOMO-silenced samples, expression of human WT NOMO1 (resistant to siRNA targeting murine NOMO1) significantly enhanced differentiation compared to non-rescued NOMO-silenced samples, though it remained somewhat below WT levels (Fig. 6c,d). We attribute this incomplete rescue to the use of a human construct in a mouse line. Next, we sought to determine whether the Ig 1/10/11 interface critical for ER morphology and force-bearing is necessary for myogenesis. To this end, we produced stably expressing F-NOMO1r_4-_^Mut^ in WT and _m_NOMO KO myoblast lines. Upon inducing differentiation under _m_NOMO knockdown, the 4-mutant-expressing cells exhibited delays in myotube formation and Myhc patterning was more pronounced than for both _m_NOMO-silenced and WT-rescued conditions (Fig. 6c,d). Together, these results suggest that NOMO acts as a myogenesis factor, dependent on critical conformational interactions and possibly load-bearing capabilities.

If NOMO is indeed required for muscle function, locomotion defects would be expected on an organismal level. To test this idea in a metazoan while avoiding viability issues arising from potential heart defects, we employed a nematode model (nematodes lack a heart). We directly compared the motility of *Caenorhabditis elegans* challenged with control RNAi with those in which the direct NOMO homolog *nra-4* was targeted via RNAi. The motility of nematodes challenged with *nra-4*-directed RNAi was significantly compromised relative to control animals (Fig. 6e). This result supports our functional assignment of NOMO to a critical role in muscle function.

## Discussion

This study establishes NOMO as a mechanosensitive protein essential for myogenesis and identifies key molecular features that are critical for force transduction and functionality. Our findings not only highlight NOMO’s involvement in muscle differentiation but also advances our understanding of how intracellular organelles and proteins process mechanical stimuli.

ER morphology is vital to cellular function and depends on a suite of protein shapers and spacers (Shibata et al., 2010; Westrate et al., 2015). We previously demonstrated that NOMO-depleted cells undergo substantial ER rearrangement (Amaya et al., 2021). The restoration of this phenotype by expression of NOMO1 or ER-shaping proteins Atlastin-2 or Climp-63, together with evidence that NOMO1 directly affects sheet spacing, established its role in maintaining ER structure. Despite these findings, the precise mechanisms by which NOMO preserves ER morphology were not fully understood. Here, we find that a highly conserved interface between Ig 1 and Ig 10-11 is central to NOMO function, as mutations disrupting these interactions severely compromise NOMO1’s ability to rescue ER morphology, buffer mechanical forces, and promote myogenesis (Fig. 2d,f and Fig. 6d,e). Our analyses furthermore demonstrate that interactions between Ig 1 and Ig 10/11 are of modest (∼1 µM) affinity (Fig 3d), yet are essential for the metastable positioning of NOMO within the ER and clearly contribute to the comparatively slow diffusional mobility (Fig. 2e,f and Fig. 4).

These findings suggest that a delicate balance between stability and dynamic flexibility may be linked to the oligomerization of NOMO through Ig domains 1, 10, and 11. We envision that NOMO may assume dynamically interchangeable and coexisting assemblies (Fig. 7) derived from the basic unit of the NOMO dimer (Fig. 4) and capable of higher-order configurations. The identical interfaces in NOMO assemblies would allow for energetically neutral switching among conformations. Lower NOMO concentrations likely favor a looping model (Fig. 1a and Fig. 7, models I and III), while higher densities promote intermolecular binding (Fig. 7, models II and IV). The ability of Ig 1-2 or Ig 10-11 to induce a dominant negative phenotype (Fig. 3f,g) suggests a dynamic interconversion of these assembly modes. This flexible equilibrium would support bulk cargo transport through the ER, maintain membrane integrity, and enable dynamic adaptation of ER morphology, including the sliding of parallel membranes in response to mechanical challenges.

**Fig. 7.**
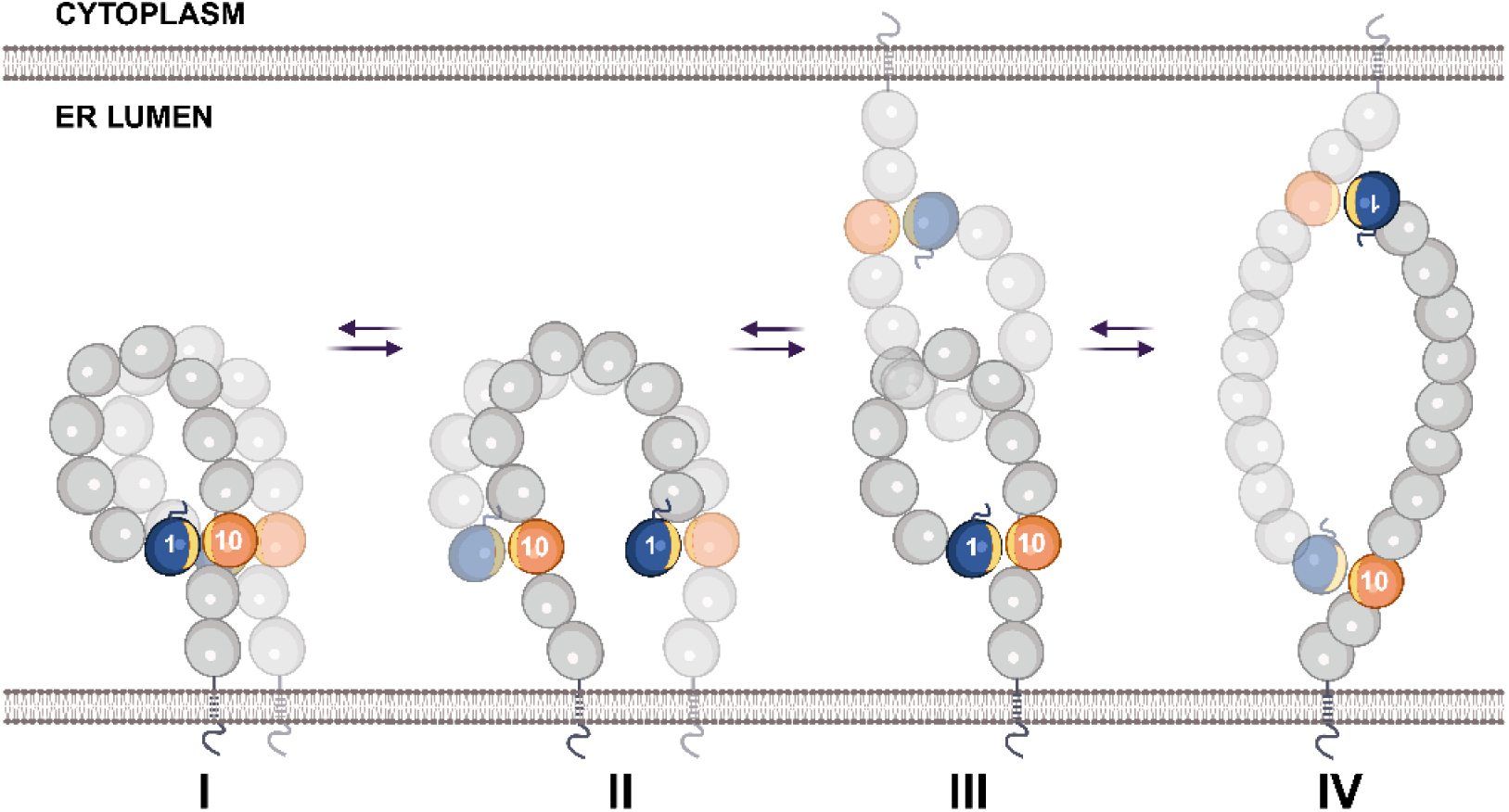
Models of NOMO assembly in the ER. NOMO dimerization is represented in cis in models I and II, and in trans in models III and IV. These models serve as the basic building blocks of NOMO oligomerization. The Ig 1/10/11 interface (yellow sliver) is shown intramolecularly in I and III and intermolecularly in II and IV. Ig domains are represented as circles, with Ig 1 and Ig 10 colored blue and orange, respectively.

Apart from homotypic assemblies, we and others have proposed that NOMO serves additional functions, e.g. by restricting other membrane protein complexes to sheet regions of the ER (Amaya et al., 2021; Lewis et al., 2024). Indeed, NOMO is an integral component of the BOS complex, where it likely anchors Nicalin and TMEM147 to regions of sheet regions where protein synthesis occurs. While not depicted, the modeled states would accommodate a NOMO-Nicalin interface at the backside of Ig 11-12 (Gemmer et al., 2024; Page et al., 2023).

The dynamic states of NOMO may be influenced by forces acting on the ER, which are essential for maintaining cellular function (Feng & Kornmann, 2018; Goyal & Blackstone, 2013; Song et al., 2024). Mechanical stimuli are transmitted through cytoskeletal interactions, membrane-associated proteins, and interactions with other organelles (McGinn et al., 2021), which initiate pathways that control cellular functions such as gene and protein expression, organelle dynamics, and cell fate decisions (Dupont & Wickstrom, 2022; Orr et al., 2006; Romani et al., 2021). Tension-sensitive probes employed in this study demonstrate that NOMO experiences mechanical stress in the range of ∼5 pN (Fig. 5), similar to that documented in other force-responsive proteins (Carley et al., 2021; Kumar et al., 2016). Sheet spacing and the Ig 1/10/11 interface modulate strain intensity (Fig. 5h and 5c,g), emphasizing the importance of conformation in mechanosensitivity (Lombardi & Lammerding, 2011). In the future, it will be interesting to test if NOMO associates with cytoskeletal components via the cytosolic tail, as was reported for Climp-63 (Klopfenstein et al., 1998). Notably, NOMO’s Ig domains closely resemble energy-absorbing regions in cadherins, titin, and pilus adhesive proteins (Echelman et al., 2016; Ren & Berro, 2022; Tskhovrebova et al., 1997; Yonemura et al., 2010) (see Supplemental Table 1 for a structural comparison), evocative of a dissipative architecture designed to handle mechanical strain. Future work will be needed to understand how dynamic forces and interactions with ER components affect NOMO’s Ig domain-mediated force buffering properties and potential roles in mechanotransduction.

The mechanosensing properties of NOMO may impact ER and sarcoplasmic reticulum (SR) morphology and broader processes such as myogenesis. We found that depleting NOMO severely impedes C2C12 myogenesis (Fig. 6d), highlighting a crucial function for NOMO in promoting muscle cell differentiation. Interestingly, NOMO is structurally similar to gp210, a nuclear pore complex (NPC) component implicated in myocyte differentiation (Gomez-Cavazos & Hetzer, 2015). Given the nuclear envelope abnormalities observed in NOMO-depleted myotubes (Fig. 6b) and earlier findings highlighting essential roles for ER and nuclear-associated proteins in muscle development and function (D’Angelo et al., 2012; Hao & Starr, 2019; Janota et al., 2020; Kalukula et al., 2022), future studies are warranted to scrutinize a possible functional interplay of NOMO, gp210 and the linker of nucleoskeleton and cytoskeleton complex (LINC) in supporting the mechanical properties of the NE (Sosa et al., 2013).

Findings from our study, along with those of others (Haffner et al., 2004), provide evidence for NOMO’s involvement in cellular differentiation processes and suggests broader implications in tissue development and homeostasis. The identification of NOMO as a force-bearing protein involved in myoblast differentiation, DM1 presentation, and nematode motility established in this study establishes its critical role in striated muscle physiology. Future research will focus on further elucidating NOMO’s mechanistic roles in relation to cellular forces and exploring its potential as a therapeutic target for muscular and cardiac disorders.

## Materials and methods

### Tissue Culture

All cells were obtained from ATCC. U2OS, MEF3T3, and HeLa cells were cultured as previously described (Amaya et al., 2021) (Prophet et al., 2022). Briefly, cells were cultured in Dulbecco’s modified Eagle’s medium supplemented with 10% (v/v) FBS (Thermo Fisher Scientific) and 100 U ml−1 of penicillin-streptomycin (Thermo Fisher Scientific).

### Immunofluorescence

Cells cultured on coverslips were washed once with PBS warmed to 37°C then fixed in 4% paraformaldehyde (+0.1% glutaraldehyde for any samples analyzing ER structure) at room temperature for 15 min and washed twice with PBS. Samples were permeabilized with 0.3% (v/v) Triton-X-100 for 3 min, washed three times with PBS for 5 min each, and blocked with 3% BSA in PBS for 1 h at room temperature. Coverslips were then placed in a humidity chamber and incubated with primary antibodies diluted in 3% BSA in PBS for 1 h at room temperature. Following three PBS washes for 5 min each, coverslips were once more placed in the humidity chamber and incubated with secondary antibodies in 3% (w/v) BSA in PBS for 1 h at room temperature in the dark. Cells were washed with PBS, incubated with Hoechst 33342 nuclear stain (Life Technologies) for 1 min, and washed twice in PBS for 5 min before being mounted onto slides using Fluoromount-G (Southern Biotech). Primary antibodies were diluted 1:500: anti-FLAG produced in mouse (F1804, Millipore Sigma), anti-FLAG produced in rabbit (D6W5B, Cell Signaling), anti-HA (11867423001, Roche) anti-calreticulin (27298-1-AP, Proteintech), anti-Myosin hc (MAB4470, R&D Systems), and diluted 1:1000 Phalloidin FITC reagent (ab235137, Abcam). Secondary antibodies were conjugated to Alexa Fluor 488 (Life Technologies) or Alexa Fluor 568 (Life Technologies).

Images were acquired on an LSM 880 laser scanning confocal microscope (Zeiss) with Airyscan using a C Plan-Apochromat 63×/1.40 oil DIC M27 objective using ZEN 2.1 software (Zeiss). Standard wide field images were obtained on a Zeiss Axio Observer D1 microscope with a 63× oil immersion objective and an AxioCam MRm microscope camera using AxioVision 4.8 software (Zeiss). For live-cell imaging, cells were grown in a µ-Slide 8 Well Glass Bottom chamber (80827; ibidi) and imaged in a CO_2_-, temperature-, and humidity-controlled Tokai Hit Stage Top Incubator. Images were acquired on an inverted Nikon Ti microscope equipped with a Yokogawa CSU-X1 confocal scanner unit with solid state 100-mW 488-nm and 50-mW 561-nm lasers, using a 60× 1.4 NA plan Apo oil immersion objective lens, and a Hamamatsu ORCA R-2 Digital CCD Camera.

### Cloning, transient RNAi knockdowns, and transient transfections

Constructs were cloned by Gibson assembly as previously described (Amaya et al., 2021) from Dharmacon plasmids containing the NOMO1 gene in a pcDNA3.1+ vector. siRNA KD experiments were executed as previously described (Rampello et al., 2020). Briefly, cells were seeded in either a 12-well plate at 0.5×10^5^ cells/well or a 6-well plate at 1×10^5^ cells/well. For KD, lipofectamine RNAimax (Invitrogen) was used to transfect 50 nM of either non-targeting or NOMO1 siRNA (Horizon Discovery), followed by a second transfection at 24 h and harvesting or fixation at 72 h. For rescues, Lipofectamine 3000 (Invitrogen) was used to transfect the siRNA/DNA mixture. Each siRNA and plasmid combination was tested in four independent experiments. Following immunofluorescence (described above), CellProfiler software (Stirling et al., 2021) was used for void and myogenesis quantification. Describe pipeline.

### Lentivirus production and myoblast transduction

Generating lentivirus for stable integration of NOMO constructs into myoblast cell lines was adapted from previously described methods (Tsai et al., 2019) and (Gomez-Cavazos & Hetzer, 2015). Low-passage HEK293T cells were seeded in 10-cm plates at 3.5 x 10^6^ cells per plate. The following day, each plate containing 6 mL of medium was co-transfected with three plasmids: MMLV gag/pol (2 μg), viral envelope protein VSV-G (1 μg), and either pBABE-puro-F-NOMO1r (6 μg) or pBABE-puro-F-NOMO1r^4-Mut^ (6 μg) using 27 µL of X-tremeGENE 9 (Roche). Supernatants were filtered after 72 h through a 0.45 μm filter unit. To proliferating myoblast cells seeded in 6-well plates at 1×10^5^ cells/well (∼30% confluency) in 2 mL medium supplemented with 4 μg/mL of polybrene, virus was serially diluted from 50-500 μL. After 48 h, 2.5 μg/mL puromycin was added to each well, including to a control well lacking transduction, and incubated for 48 h. Cells were seeded in 15-cm plates at low density (∼150 cells/plate) and isolated single clonal clusters were validated by immunofluorescence.

### Immunoblot analysis

For protein extracts, cells were lysed in 1% SDS buffer with benzonase and incubated at 95°C for 5 min. For analysis of C2C12 attached and detached populations, detached cells were harvested by collecting the floating and loosely attached cells in the media and recovered living cells by scraping the plates. Protein concentration was determined using the bicinchoninic acid reagent (Thermo Fisher Scientific). For immunoblot blot analysis, equal amounts of protein (10-20 µg) were resolved by 8% SDS–PAGE gels and transferred onto polyvinylidene fluoride membranes (Bio-Rad). The membranes were blocked in 4% (wt/vol) non-fat milk in PBS + 0.1% (vol/vol) Tween-20 (Sigma-Aldrich). Primary and horseradish peroxidase-conjugated secondary antibodies were diluted in blocking buffer. The blots were visualized by chemiluminescence on a ChemiDoc gel imaging system (Bio-Rad).

### FRAP

FRAP experiments were performed using an LSM 880 laser scanning confocal microscope (Zeiss) using a C Plan-Apochromat 63×/1.40 oil DIC M27 objective using ZEN 2.1 software (Zeiss). Experiments were done at 37°C with 5% CO_2_ using a live-cell chamber system. For each acquisition, NOMO-TS_in_ was bleached using the 488 nm laser. The eGFP in TS was subjected to photobleaching, leaving tagRFP unperturbed. Between five pre-bleach images were acquired and post-bleach images were acquired every 0.15 s for 60 s. To monitor recovery, a circular ROI of 1 µm was used to measure the average fluorescence intensity before and after photobleaching. These values were normalized to the pre-bleach frame and graphed relative to bleaching. Data from at least three experiments totaling at least fifty cells for each construct were collected and the averaged FRAP measurements were plotted in Graphpad Prism 10 and fit to a nonlinear regression exponential curve. A GFP-NOMO1 construct served as a control reference to confirm the tension sensor module did not interfere with NOMO dynamics and recovery time (data not shown).

### FRET

The eGFP-TagRFP tension sensor module (TS) (kind gift from Martin Schwartz) was inserted after NOMO Ig 12 and prior to the transmembrane domain. The no-tension (NOMO^LD^-TS) construct was generated by inserting a stop codon after the sensor before the transmembrane domain. Cells were plated on glass-bottomed dishes (CELLTREAT) coated with poly-L-lysine (PLL) (Sigma P9155) and transfected using Lipofectamine 2000 or Lipofectamine 3000 reagent (ThermoFisher) according to the manufacturer’s instructions and imaged the following day. Untransfected cells were used as dark controls. Imaging was performed on a LSM 880 laser scanning confocal microscope (Zeiss) using a C Plan-Apochromat 63×/1.40 oil DIC M27 objective using ZEN 2.1 software (Zeiss) with a stage maintained at 37°C and 5% CO_2_. Images were acquired using Zen software. FRET imaging was performed as previously described (Kumar et al., 2016) (Grashoff et al., 2010). Briefly, a 0.5-1 µm ROI in the cellular midplane of the ER was imaged five to ten times for eGFP (ex. 488-nm laser, em. 527/55 filter) and TagRFP (ex. 561 nm laser, em. 615/70 nm filter) prior to acceptor photobleaching using the 561-nm laser line. Five to ten sequential images were acquired for both donor and acceptor and the normalized FRET index was calculated by taking the ratio of donor intensity images before and after bleaching the acceptor: 1 – I*_donor pre bleach_*//I*_donor post bleach_*.

### Recombinant protein expression and purification

For NOMO Ig expression, N-terminally His_6_-tagged Ig 1-2, Ig 1-2^3-Mut^, and Ig 10-11 proteins were each expressed in BL21(DE3) *E. coli* strains. 4 L cultures were grown in terrific broth (TB) at 37 °C with shaking and expression was induced at an OD_600_ of 0.7 with 1 mM isopropyl β-D-1-thiogalactopyranoside (IPTG) for 4 h at 20 °C. Cells were centrifuged at 8,000xg for 30 min at 4°C frozen at −80°C until use. Pellets were resuspended in lysis buffer (50 mM Tris pH 8, 50 mM NaCl, 2 mM 2-Mercaptoethanol (BME), 10 mM imidazole, 1x Roche cOmplate EDTA-free Protease Inhibitor Cocktail (PI), 30 µM PMSF, 10% (v/v) glycerol), then subjected to three iterations of French pressure cell press. Following centrifugation at 20,000xg for 30 min at 4 °C, supernatants were incubated with Ni-NTA agarose (Qiagen) for two hours with gentle shaking at 4 °C. Samples were run through gravity flow columns and washed with 50 mL of wash buffer (50 mM Tris pH 7.5, 300 mM NaCl, 5 mM MgCl_2_, 2 mM βME, 1x PI, 25 mM imidazole). To elute, 5 mL of elution buffer (50 mM Tris pH 7.5, 150 mM NaCl, 2 mM MgCl_2_, 1 mM βME, 250 mM imidazole) was added to the column and collected samples were assessed for purity by SDS-PAGE. Eluates were dialyzed overnight in 50 mM Tris pH 7.5,150 mM NaCl, and 1 mM βME and then subjected to size-exclusion chromatography with an S75 column. Samples were concentrated at 500xg when needed using Amicon® Ultra Centrifugal Filters.

For full-length NOMO expression, Expi293F cells were transfected with either Flag-NOMO^WT^ or Flag-NOMO^4-Mut^ constructs using the ExpiFectamine 293 Transfection Kit (Gibco) following the manufacturer’s protocol for a 100 mL culture. Cells were harvested 72 h later and lysed in 60 mL of lysis buffer (50 mM MES pH 6.0, 100 mM NaCl, 50 mM KCl, 5 mM CaCl_2_, 5% glycerol, 1% n-Dodecyl-β-D-maltoside (DDM)) for 1 h at 4°C. Lysates were spun for 30 min at 20,000xg at 4°C and the resulting supernatants incubated with 0.5 mL anti-FLAG M2 beads (Sigma) overnight. Extracts were then loaded onto a gravity column and washed with 50 mL of buffer (50 mM MES pH 6.0, 150 mM NaCl, 100 mM KCl, 5 mM CaCl_2_, 2% glycerol, 0.05% DDM). To elute FLAG-tagged NOMO, beads were incubated for 30 min with 2 mL of elution buffer (50 mM MES pH 6.0, 150 mM KCl, 5 mM CaCl_2_, 5 mM MgCl_2_, 2% glycerol, 0.05% DDM, 15 μM FLAG peptide). The elution was concentrated at 100xg to 10 µM and subjected to SEC in an S200 or S75 column (GE Healthcare).

### SEC and SEC-MALS

For *in vitro* binding studies, His_6_-Ig 10-11 was incubated with either His_6_-Ig 1-2 or Ig 1-2^3-Mut^ in buffer containing 50 mM Tris-HCl pH 7.5, 100 mM NaCl, 5 mM MgCl_2_, and 5 mM CaCl_2_. The mixture was incubated at room temperature for one hour prior to running on an equilibriated Analytical Superdex-75 10/300 (Cytiva). His_6_-Ig 10-11 and His_6_-Ig 1-2^3-Mut^ binding was tested with an overnight incubation, as well as a 1:2 ratio of His_6_-Ig 10-11 and His_6_-1-2^3-Mut^. Apparent molecular masses were determined by calibrating the size-exclusion column with the following set of protein standards: thyroglobulin, ferritin, catalase, aldolase, albumin, ovalbumin, chymotrypsinogen, and ribonuclease A.

Multi-angle laser light-scattering experiments were performed at room temperature in 50 mM MES pH 6.0, 150 mM KCl, 5 mM CaCl_2_, 5 mM MgCl_2_, 2% glycerol, and 0.05% DDM. Light-scattering data were measured using a Dawn Heleos-II spectrometer (Wyatt Technology) coupled to an Opti-lab T-rEX (Wyatt Technologies) interferometric refractometer. Prior to sample runs, the system was calibrated and normalized using the monomeric bovine serum albumin as an isotropic protein standard. Samples at 1-1.5 mg/mL in 250 µL were injected into a Superdex 200 Increase 10/300 GL column (GE Healthcare) at a 0.5 ml/min flow rate. Data on light scattering (690 nm laser), UV absorbance (280 nm), and refractive index were measured simultaneously during the run. Data were processed in ASTRA software.

### Isothermal Titration Calorimetry Assay

ITC measurements were executed on a VP-ITC Microcal calorimeter (Microcal) at 25°C. Buffer contained 50 mM Tris-HCl pH 7.5, 100 mM NaCl, 5 mM MgCl_2_, and 5 mM CaCl_2_. All solutions were filtered and degassed. Starting with an upper cell analyte concentration of 300 µM His_6_-Ig 10-11 – above which the fragment precipitated – we adjusted the titrant concentration to 20 µM for His_6_-Ig 1-2 and His_6_-Ig 1-2^3-Mut^. This minimum value ensured sufficient heat detection above the background, which was necessary due to the smaller enthalpic contribution compared to the entropic contribution of the interaction. Automated titrations injected 10-15 μl of His_6_-Ig 10-11 from a syringe into the sample cell containing either buffer, His_6_-Ig 1-2, or His_6_-Ig 1-2^3-Mut^ at a time interval of 120 s and repeated 22-33 times. The heat of interaction between both of the components was measured after subtracting the heat of dilution and baseline was subtracted. The titration data were analyzed by Origin7.0 (Microcal).

### Sequencing data

RNA sequencing data for control and DM1 Tibialis anterior muscle biopsies is publicly available (GSE86356) and was published by (Wang et al. 2019: Transcriptome alterations in myotonic dystrophy skeletal muscle and heart). Raw reads were processed as previously described (Todorow et al., 2022). Normalized counts for NOMO1 were plotted against decreasing dorsiflexion strength as assessed in (Wang et al. 2019: Transcriptome alterations in myotonic dystrophy skeletal muscle and heart).

### Myoblast differentiation assays

C2C12 myoblast cells differentiation into myotubes was induced by shifting confluent (∼80%) myoblasts from DMEM containing 10% FBS to media containing 2% (v/v) horse serum. Stable cell lines were generated in independent triplicates by lentiviral transduction and consisted of mixed populations. For differentiation experiments, cell lines were plated in 6-well tissue-cultured treated plates (Corning) at the same density and induced to differentiate upon confluency. Fixation and immunostaining were performed as described above. Cellprofiler was used to quantify differentiation, whereby the number of nuclei per image were binned according to whether they were present in an Myhc+ myotube.

### *C. elegans* experiments

Nematodes expressing *myo-3p::myo-3::gfp* were maintained at 25 °C. RNAi against nicotinic acid receptor protein-4 (*nra-4*) was induced through feeding of *E. coli* HT115(DE3) expressing *nra-4* dsRNA obtained from the Ahringer RNAi library (Kamath & Ahringer, 2003). Nematodes were fed RNAi-containing bacteria and its respective control (HT115(DE3) expressing empty vector L4440) for two generations prior to the motility analysis. Nematodes were synchronized manually by egg laying. The motility of day 4 old nematodes (young adults) was analyzed in M9-media. Nematodes were dispersed in fresh M9-media (750 µl in a 3.5 cm culture plate) and left to acclimatize for 30 seconds before video acquisition at 7.8-fold magnification and 25 frames per second. The videos were recorded in bright field using a Leica stereo microscope M205FA. 5-7 nematodes were recorded at once and a total of at least 15-20 nematodes were recorded per replicate and per condition. Videos were analyzed using WormLab software (WormLab 2023.1.1 (MBF Bioscience LLC, Williston, VT USA)). Body bends per second were calculated through the body angle-midpoint function of the software. Overlapping nematodes were censored from the analysis. Statistical analysis (one way ANOVA) was performed using GraphPad Prism. P-values were considered significant when: p ≤ 0.05 (*), p ≤ 0.01 (**), p ≤ 0.001 (***).

---

**Extended Data Fig. 1.**
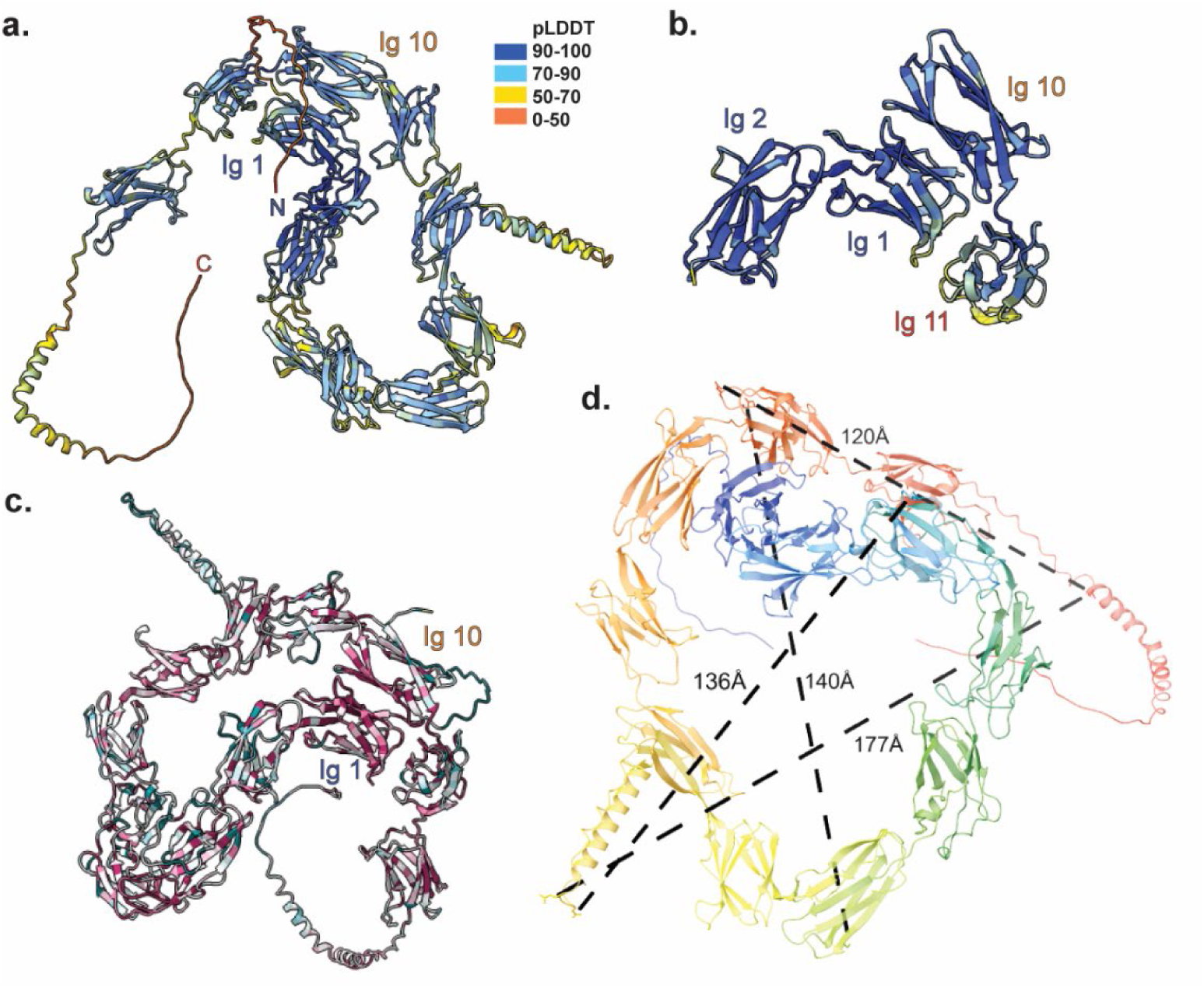
NOMO structure predictions are confident and conserved. a, Alphafold3 models a highly confident structure of NOMO, colored by the per-residue confidence metric predicted local distance difference test (pLDDT) on a scale from 0 to 100, depicted in the legend inset. b, Ig 1-2 and Ig 10-11 colored by pLDDT. c, NOMO FL protein sequence conservation for as predicted by ConSurf. Color scale same as in Fig. 1e. d, Representative distances in angstroms (Å) between distal regions in NOMO.

**Extended Data Fig. 2.**
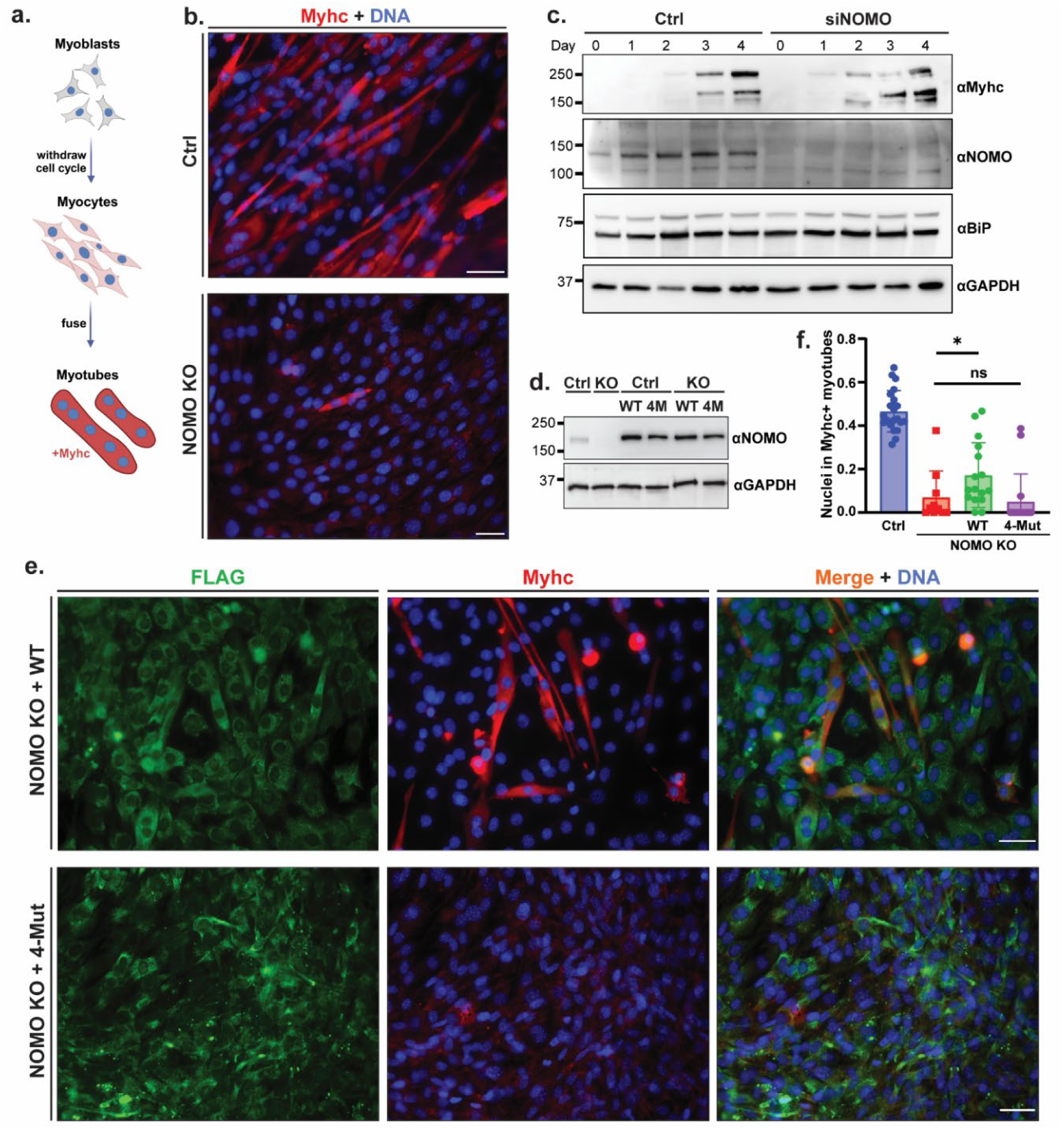
C2C12 differentiation in NOMO knockout cells. a, Cartoon representation of C2C12 myogenesis, whereby myoblasts exit the cell cycle to differentiate into myocytes, which then fuse to form multi-nucleated myotubes, driving muscle fiber maturation. b, Myogenesis performed as in Fig. 6a and immunostained for Myhc and stained for DNA with DAPI. c, Immunoblot of control or NOMO silenced (siNOMO) myoblasts during indicated days of differentiation. d, Immunoblot of indicated NOMO rescue constructs in either WT (Ctrl) or NOMO knockout (KO) myoblast lines. e, Assay performed as in b, with myoblasts stably expressing either WT (F-NOMO^WT^) or 4-Mut (F-NOMO^4-Mut^) NOMO1 constructs in NOMO knockout lines. f, Quantification of myogenesis under indicated conditions, scored by the fraction of nuclei in Myhc+ myotubes. Statistical analyses were performed using a two-tailed unpaired Mann– Whitney test; *P < 0.05 and ns, not signficiant. Error bars show ± s.e.m. All scale bars, 50 μm. Myhc, myosin heavy chain.

**Extended Data Fig. 3.**
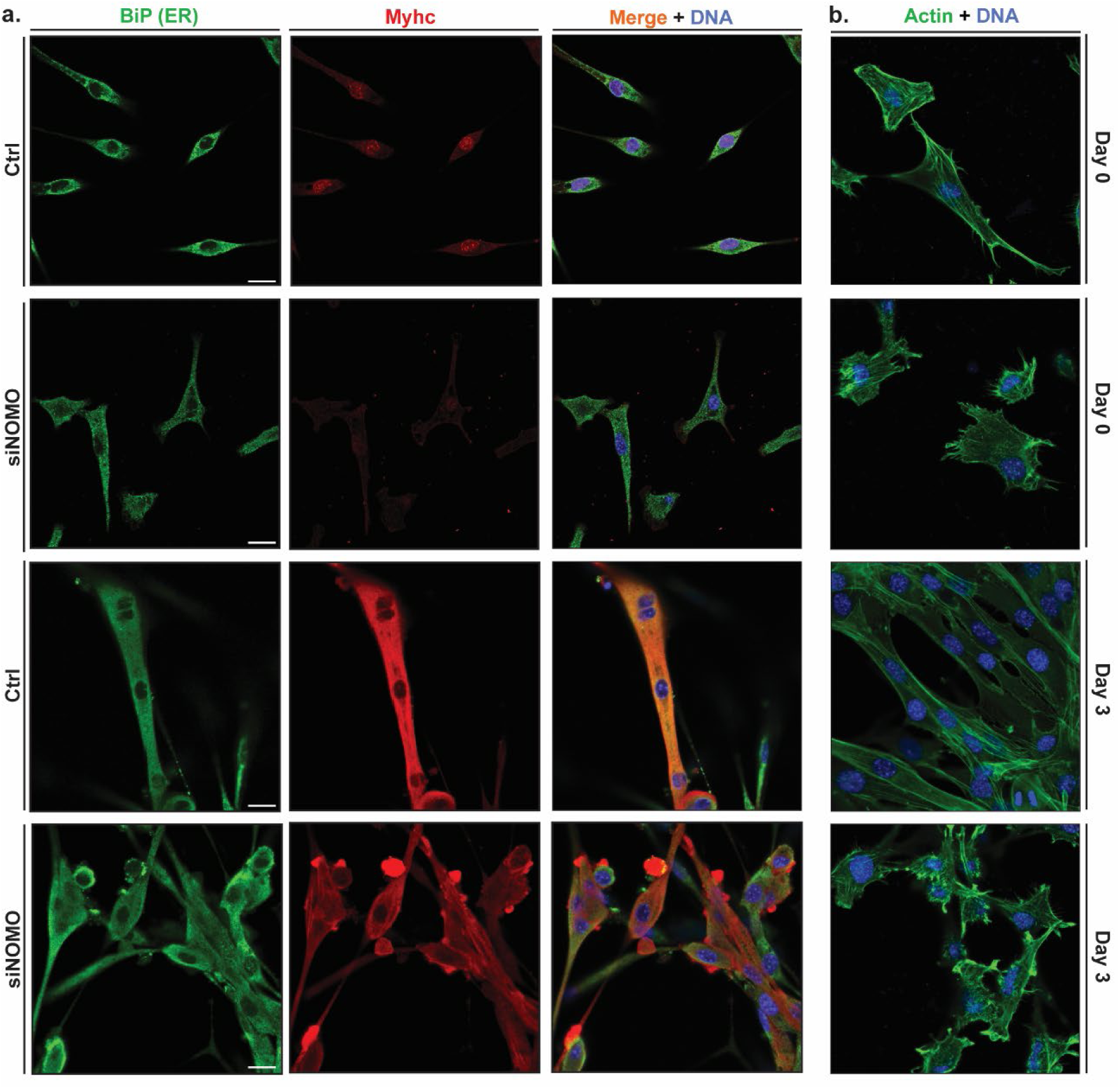
ER, actin, and Myhc morphology effects under NOMO knockdown. a, Myoblast cells pre-differentiation (Day 0) and post-differentiation (Day 3) into myocytes and myotubes, immunostained for Myhc and ER marker BiP. b, Conditions as in a, probed for actin by FITC-phalloidin. Myhc, myosin heavy chain.

**Extended Data Fig. 4.**
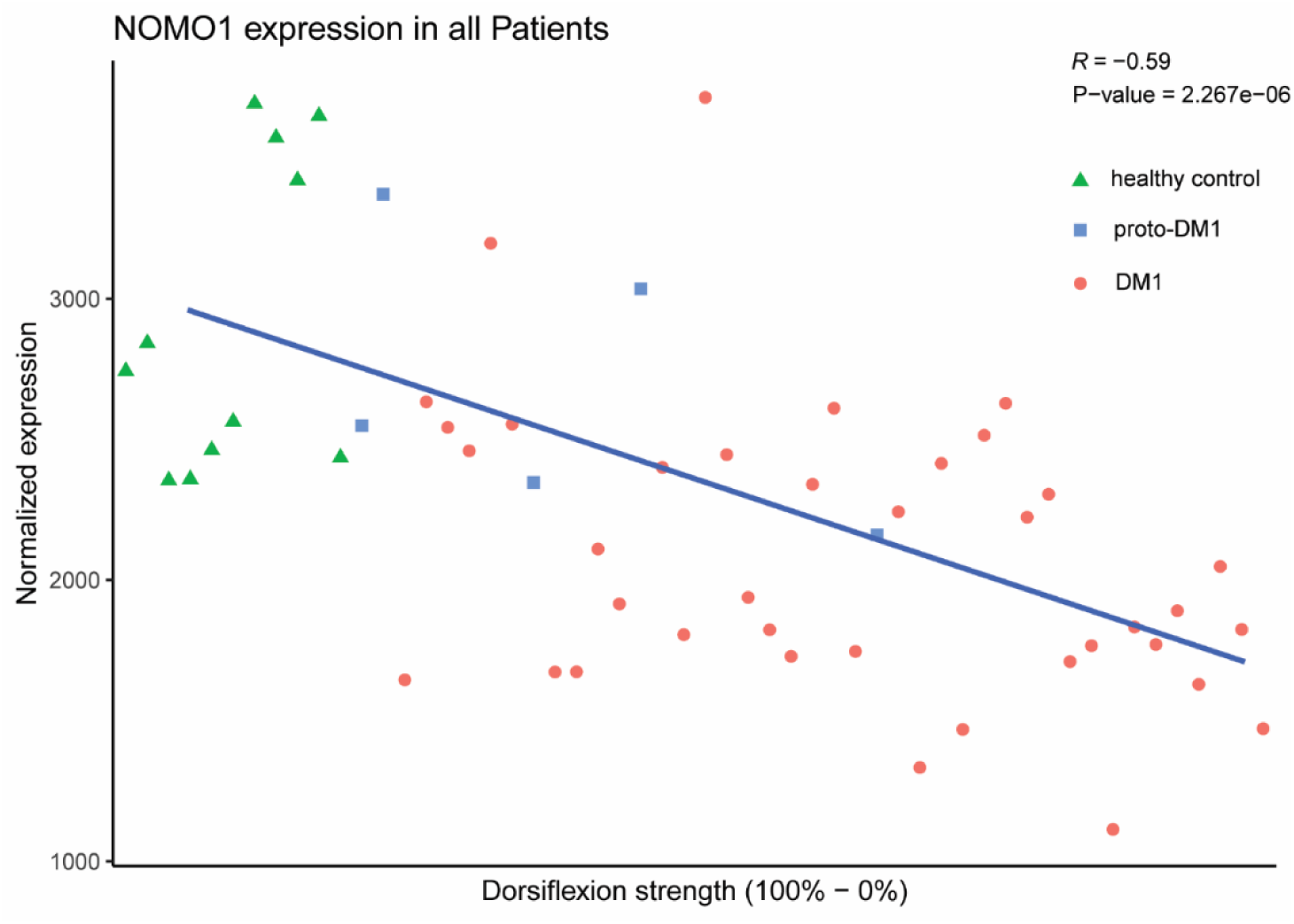
Normalized expression of NOMO1 in T. anterior muscle biopsies. Dorsiflexion strength measured in control and DM1 patients against NOMO transcript levels (R = −0.59). DM1, Myotonic dystrophy type 1. proto-DM1, intermediate form of DM1 corresponding to fewer CTG-repeats (50-100) than DM1 (>100) or healthy controls (<30).

**Supplemental Table 1:** Dali structure comparison.

| PBD - chain | Entry | Gene | Z-score | Description |
| --- | --- | --- | --- | --- |
| 3rkp | Q81D71 | Q81D71_BACCR | 11.8 | Collagen Adhesion Protein |
| 8bdw-H | F9UR90 | lp_2578 | 11.5 | Cell Surface Adherence Protein,Collagen-Binding |
| 5aq0 | O75976 | CBPD_HUMAN | 11.4 | Carboxypeptidase D |
| 3htl-X | Q6NF81 | DIP2013 | 11.3 | Putative Surface-Anchored Fimbrial Subunit |
| 1qmu | P83852 | CBPD_LOPSP | 11.2 | Carboxypeptidase Gp180 Residues 503-882 |
| 8vdk |  |  | 11.2 | Serine-Aspartate Repeat-Containing Protein D |
| 4oq1-A | A0A0H2UNM0 | SP_0464 | 11.1 | Cell Wall Surface Anchor Family Protein |
| 3kpt-B | Q81D71 | BC_2508 | 11 | Collagen Adhesion Protein |
| 3phs-A | Q8E0S8 | SAG0646 | 10.8 | Cell Wall Surface Anchor Family Protein |
| 4jdz |  |  | 10.5 | Ser-Asp Rich Fibrinogen/Bone Sialoprotein-Binding |
| 4p0d-A | P18481 | tee6 M6_Spy0160 | 10.3 | Trypsin-Resistant Surface T6 Protein |
| 5hts-A |  | LRHM_0426 | 10 | Cell Surface Protein SpaA |
| 4eiu-A | A7V680 | BACUNI_03093 | 10 | Uncharacterized Hypothetical Protein |
| 1r6v-A | Q93LQ6 | fls | 9.9 | Subtilisin-Like Serine Protease |
| 6fwv-A | A0A6L8P957 | GBAA_5258 | 9.9 | Collagen Adhesion Protein |
| 5f44 |  |  | 9.9 | Cell Surface Protein Spaa |
| 5j4m |  |  | 9.8 | Cell Surface Protein Spaa |
| 5nv4 | G0SB58 | G0SB58_CHATD | 9.8 | Udp-Glucose-Glycoprotein Glucosyltransferase-Like |
| 5n2j-B | G0SB58 | CTHT_0048990 | 9.6 | Udp-Glucose-Glycoprotein Glucosyltransferase-Like |
| 4uzg-A | A0A0M3KKY8 |  | 9.6 | Surface Protein Spb1 |
| 6trf-A | G0SB58 | CTHT_0048990 | 9.5 | Udp-Glucose-Glycoprotein Glucosyltransferase-Like |
| 3irp-X | Q4A0V8 | uafA SSP0135 | 9.5 | Uro-Adherence Factor A |
| 2ww8-A | A0A0H2UNT6 | SP_0462 | 9.1 | Cell Wall Surface Anchor Family Protein |
| 2x5p-A | Q8G9G1 | fba2 E0F66_06355 | 9.1 | Fibronectin Binding Protein |
| 6ts2-D | G0SB58 | CTHT_0048990 | 8.9 | Udp-Glucose-Glycoprotein Glucosyltransferase-Like |
| 5m0y-B | A3DGE8 | Cthe_1806 | 8.9 | Ig Domain Protein Group 2 Domain Protein |
| 7cbs-A |  |  | 8.8 | Lpxtg Cell Wall Anchor Domain-Containing Protein |
| 2y1v-A | A0A0H2UNM7 | SP_0463 | 8.7 | Cell Wall Surface Anchor Family Protein |
| 3uxf-A | P18477 |  | 8.7 | Fimbrial Subunit Type 1 |
| 4hss-A | Q6NK05 | DIP0235 | 8.7 | Putative Fimbrial Subunit |
| 1h8l-A | P83852 | CPD | 8.7 | Carboxypeptidase Gp180 Residues 503-882 |
| 3is0 | Q4A0V8 | UAFA_STAS1 | 8.7 | Uro-Adherence Factor A |
| 8kcl |  |  | 8.7 | Lpxtg-Motif Cell Wall Anchor Domain Protein |
| 2nsm-A | P15169 | CPN1 ACBP | 8.6 | Carboxypeptidase N Catalytic Chain |
| 3irz | Q4A0V8 | Q0SWL8_CLOPS | 8.6 | Uro-Adherence Factor A |
| 6ixy | Q0SWL8 | A0A0H2UNM7_STRPN | 8.6 | Pilin |
| 8wb8 |  |  | 8.6 | Lpxtg-Motif Cell Wall Anchor Domain Protein |
| 6ts8-A | G0SB58 | CTHT_0048990 | 8.5 | Udp-Glucose-Glycoprotein Glucosyltransferase-Like |
| 7woi-A |  |  | 8.5 | Spa2 |
| 5yu5 |  |  | 8.5 | Pilus Assembly Protein |
| 2xtl |  |  | 8.5 | Cell Wall Surface Anchor Family Protein |
| 2x9x | A0A0H2UNM7 |  | 8.5 | Cell Wall Surface Anchor Family Protein |
| 3mn8-B | P42787 | svr CPD CpepE CG4122 | 8.4 | Lp15968P |
| 8j0o-G | O43402 | EMC8 C16orf2 C16orf4 | 8.4 | ER Membrane Protein Complex Subunit 1 |
| 3is1 | Q4A0V8 | UAFA_STAS1 | 8.4 | Uro-Adherence Factor A |
| 5fie | Q06853 |  | 8.4 | Cell Surface Protein Spaa |
| 5k39 | A3DGE8 | SLAP2_ACET2 | 8.3 | Cellulosome Anchoring Protein Cohesin Region |
| 5hdl |  |  | 8.2 | Cell Surface Protein Spaa |
| 5z0z |  |  | 8.2 | Pilus Assembly Protein |

## Acknowledgements

This work is supported by DOD PR200788 (C.S.), a Bachmann-Strauss Dystonia Fellowship (S.M.), and the Dystonia Medical Research Foundation (C.S.). We thank S. Bahmanyar and members of her lab for help with imaging and M. Schwartz and A. Kumar for sharing constructs and providing guidance. We thank the Yale Biophysical Resource Laboratory and Morven Graham of the Yale Center for Cellular and Molecular Imaging for assistance with electron microscopy.

## Author contributions

Conceptualization, BSN and CS; methodology, BSN, SCD, SM, SJ, YR, VT, JB, JK, YX, and CS; validation, BSN, SM, SJ, JK, and CS; formal analysis, BSN, SCD and CS; investigation, BSN, and CS; resources, BSN and CS; writing—original draft preparation, BSN.; writing—review and editing, BSN, and CS; visualization, BSN and CS; supervision, CS; funding acquisition, CS. All authors have read and agreed to the published version of the manuscript.

